# Seasonal influenza circulation patterns and projections for 2017-2018

**DOI:** 10.1101/113035

**Authors:** Trevor Bedford, Richard A. Neher

## Abstract

This is not meant as a comprehensive report of recent influenza evolution, but is instead intended as particular observations that may be of relevance. Please also note that observed patterns reflect the GISAID database and may not be entirely representative of underlying dynamics. All analyses are based on the nextflu pipeline [1] with continual updates posted to nextflu.org. We arrive at the following results:

**H3N2:** In H3N2, clade 3c2.a has continued to diversify genetically with complicated and rapid dynamics of different subclades. This diversification is not reflected in serological data that shows only minor to moderate antigenic evolution. Nevertheless, the highly parallel mutation patterns and the rapid rise and fall of clades suggests competitive dynamics of phenotypically distinct viruses.

**H1N1pdm:** Very few H1N1pdm viruses have been observed in recent months. The dominant clade continues to be 6b.1 and there is little amino acid sequence variation within HA. The only notable subclade that has been growing recently is the clade bearing HA1:R205K/S183P. This clade is dominated by North American viruses and we see no evidence that this clade has a particular competitive advantage.

**B/Vic:** Clade 1A has continued to dominate and mutation 117V has all but taken over the global population. The rise of this mutation was fairly gradual and we have no evidence that it is associated with antigenic change or other benefit to the virus.

**B/Yam:** Clade 3 has continued to dominate. Within clade 3, a clade with mutation HA1:251V is globally at frequency of about 80% throughout 2016. Within this clade, mutation 211R is at 25% frequency. In addition, a clade without prominent amino acid mutations has been rising throughout 2016.

## A/H3N2

**In H3N2, clade 3c2.a has continued to diversify genetically with complicated and rapid dynamics of different subclades. This diversification is not reflected in serological data that shows only minor to moderate antigenic evolution. Nevertheless, the highly parallel mutation patterns and the rapid rise and fall of clades suggests competitive dynamics of phenotypically distinct viruses.**

We base our primary analysis on a set of viruses collected between Jan 2015 and Feb 2017 (Fig. 1). We analyze mutation frequency trajectories in different geographic regions using all available sequence data for each region. Our phylogenetic analysis is based on a subsample of approximately 100 viruses per month where available and seeking to equilibrate sample counts geographically where possible. This equilibration attempts to collect equal samples from Africa, China, Europe, Japan/South Korea, North America, Oceania, South America, South Asia, Southeast Asia and West Asia. In the following analysis we collapse samples from China, South Asia, Southeast Asia, Japan and Korea into a single region referred to here as “Asia”, resulting in Asia possessing greater sample counts than North America or Europe. There is reasonably broad geographic sampling throughout, except Dec 2016 and Jan 2017, which have more samples from Europe and North America.

**Figure 1.**
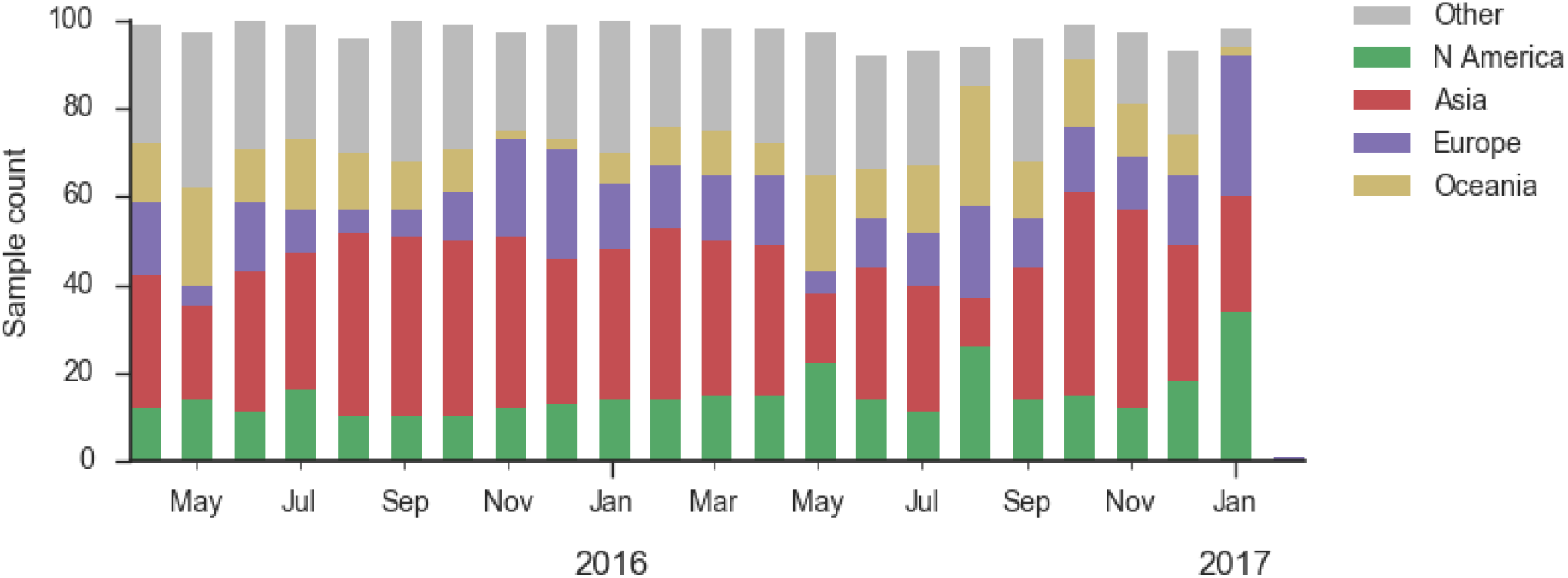
Sample counts through time and across regions. This is a stacked bar plot, so that most months there are ~100 total samples and ~15 samples each from North America and from Europe.

Of the clades that emerged from the Texas/2012 background in early 2014, only 3c2.a currently remains at globally high prevalence (Fig. 2). 3c3.b viruses have scarcly been seen since the end of 2015. The uptick of 3c3.a viruses in mid-2016 in the US has failed to carry forward to the 2016-2017 US winter season. Oceania still harbors 3c3.a viruses, but we fully expect these to be replaced by another clade in the coming year as Oceania is reseeded from the global reservoir.

**Figure 2.**
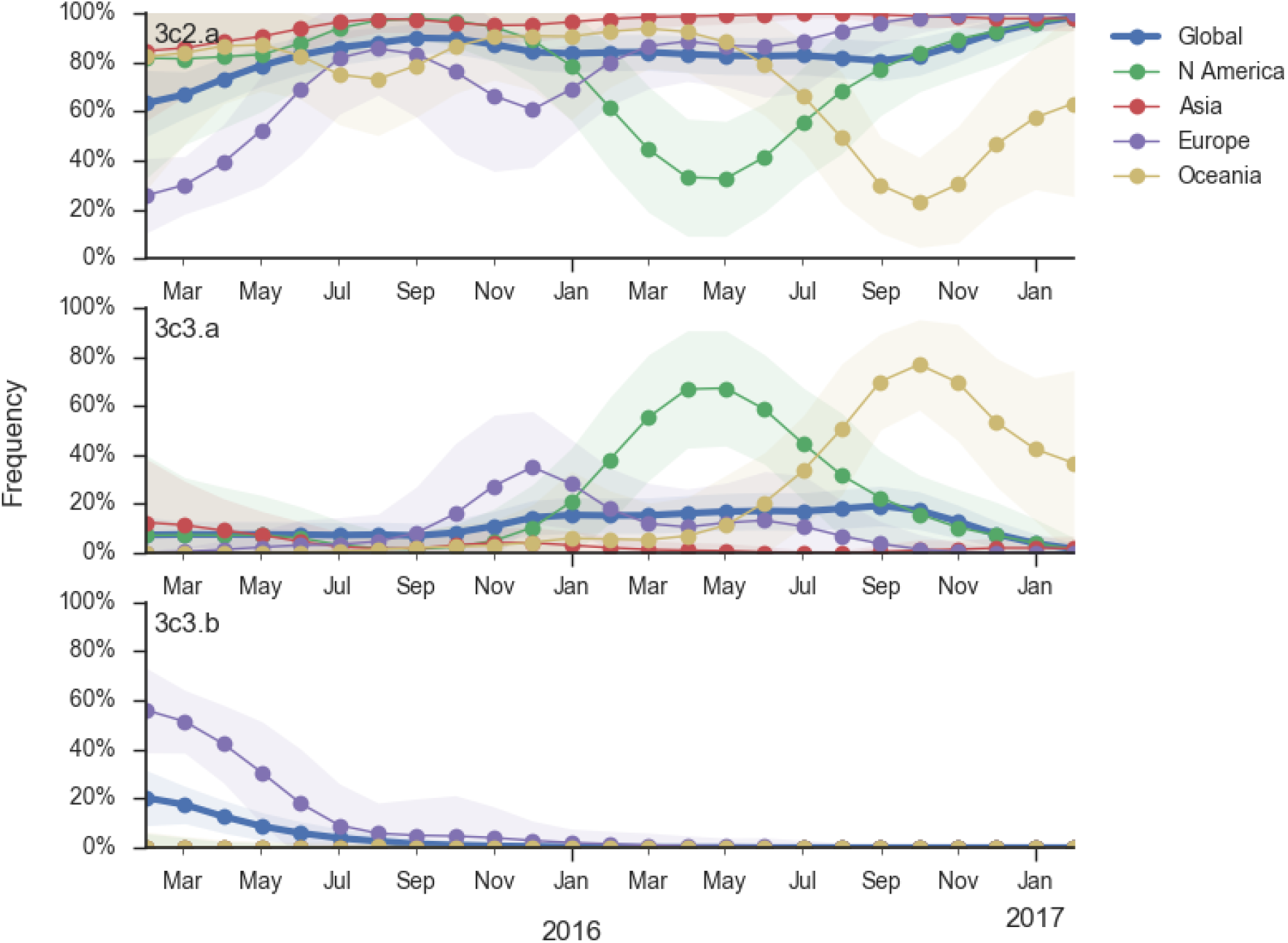
Frequency trajectories of H3N2 clades. We estimate frequencies of different clades based on sample counts and collection dates. We use a Brownian motion process prior to smooth frequencies from month-to-month. Transparent bands show an estimate the 95% confidence interval based on sample counts. The final point represents our frequency estimate for Feb 1 2017.

Within clade 3c2.a, a number of subclades have emerged (Fig. 3). Notable clades include N171K (clade 3c2a.1), N171K/N121K, N121K/S144K, 131K/142K and R142K/Q197K. Clades with R142K, N121K/R144K, Q197K arose in late 2014 or early 2015 but have not come to dominate. However, the 171K clade (3c2a.1) that was first observed in mid-2015 has continued to increase in frequency at a moderate rate and is now at a global frequency of approximately 60%. Within the 171K clade, a subclade bearing the 121K mutation emerged in at the beginning of 2016 and has steadily increased in frequency now making of the majority of the parent 171K clade.

**Figure 3.**
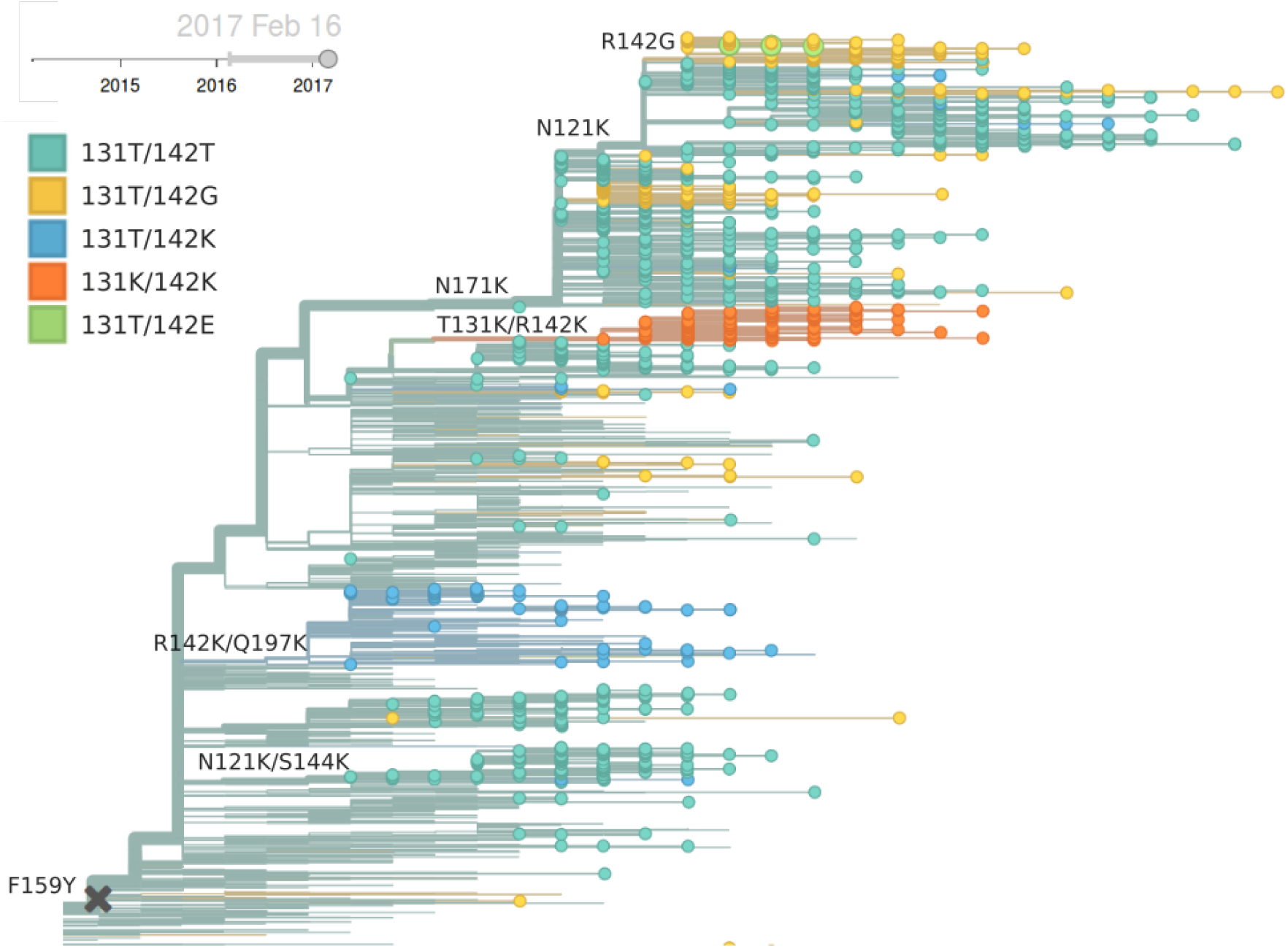
H3N2 / 3c2.a phylogeny colored by genotype.

Currently, the only noteworthy sister clade of 171K is the clade with mutations T131K/R142K (Fig. 4). This clade was first observed mid-2016 in China and dominated in China during last 3 month according to sequences available in GISAID, making up 80% of Chinese isolate since October. It has been rising rapidly and has spread to Europe and North America. In the branch leading up to this clade, we observe a rapid succession of mutation in codon 131 from threonine to asparagine and then to lysine and the mutation R142K. Rapid succession of multiple mutations in the same codon suggest that these changes are adaptive. Interestingly, we observe a large number of mutations to positively charged residues and particularly many parallel mutations to lysine at positions 121, 131, 142, 144, 171, and 197. Clades N171K/N121K and T131K/R142K are on opposing genetic backgrounds so that there is competition between these clades; both cannot succeed.

**Figure 4.**
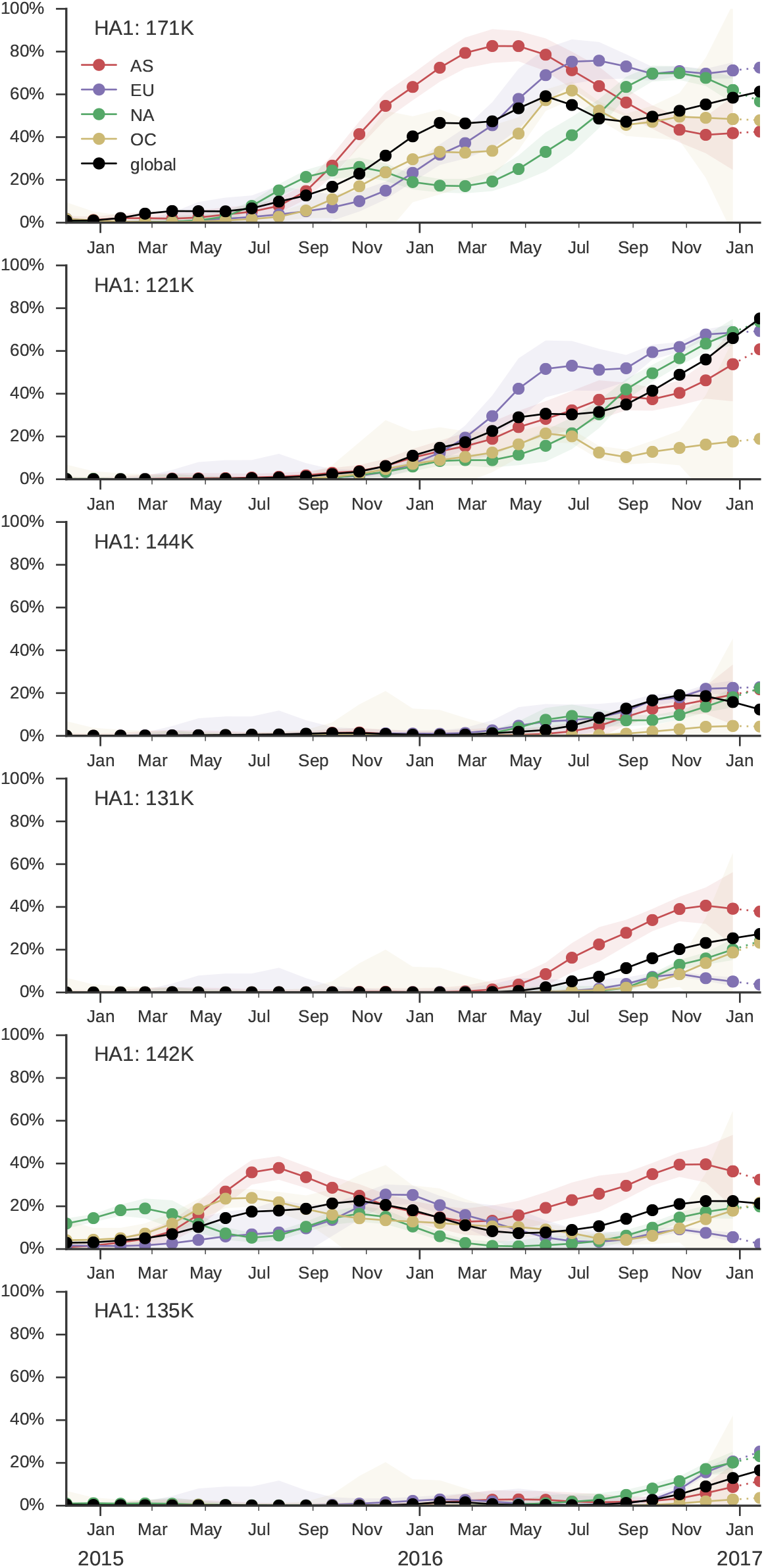
Frequency trajectories of 3c2.a subclades. We estimate frequencies of different clades based on sample counts and collection dates. We use a Brownian motion process prior to smooth frequencies from month-to-month. Transparent bands show an estimate the 95% confidence interval based on sample counts. The final point represents our frequency estimate for Feb 1 2017.

Despite the large genetic diversity at sites in the vicinity of the receptor binding site, HI or FRA assays show no evidence of substantial antigenic change (Fig. 5). HI assays by the WHO collaborating center at the Crick Institute in London indicate a moderate two-fold titer drop relative to A/HongKong/4801/2014 for several subclades, including the 171K subclade. However, there is substantial variability in those data. More dense and recent data by the CDC in Atlanta, Georgia, do not support a homogeneous titer drop of the 171K clade relative to the current vaccine strain Hong Kong/4801/2014. The figure below shows trees colored by predicted titer drop relative to Hong Kong/4801/2014 using the tree model [2] trained on HI data from Crick and CDC as well as FRA data from CDC. Taken together, these data provide little evidence for substantial antigenic evolution within 3c2.a.

**Figure 5.**
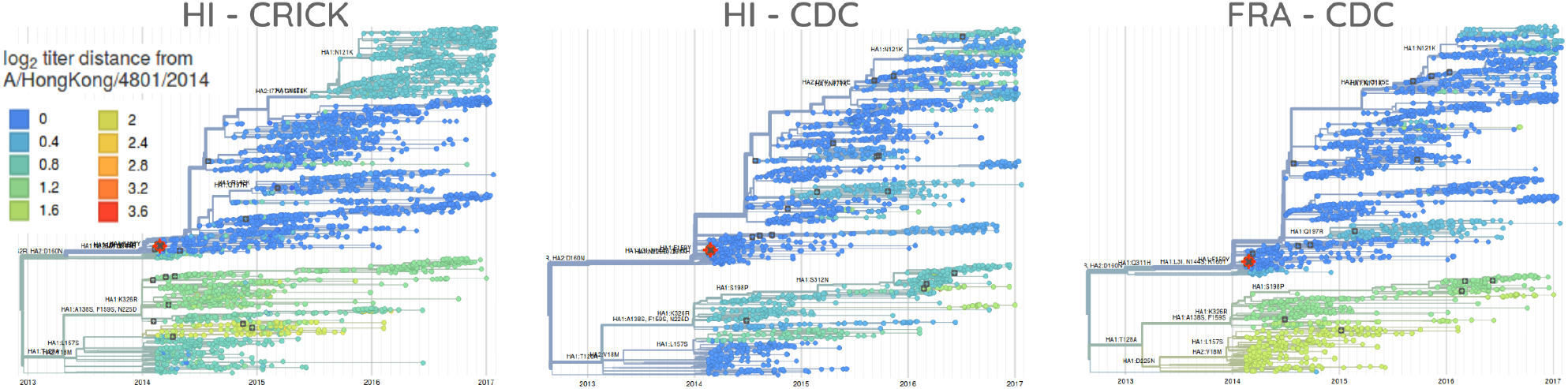
H3N2 phylogeny colored by antigenicity. Each panel shows model estimates of antigenic divergence relative to serum from A/HongKong/5738/2014. Cooler color indicates greater antigenic similarity (less titer drop going from homologous to heterologous titers). These estimates combine HI measurements from many different sera. The left panel uses HI data from the Crick Institute in London, the center panel uses HI data from the CDC in Atlanta and the right panel uses FRA data from the CDC. Crick data is current through Sep 2016 and CDC data is current through Feb 2017.

The local branching index (LBI), a phylogenetic indicator of clade growth [3], corroborates the observations made on the basis of clade frequencies (Fig. 6). Over the last 6 monthes, the 131K clade has highest LBI, while the clade 171K has highest LBI when considering sequences in all of 2016. Interpretation of the sequence based indices such as number of epitope or non-epitope mutations [4] is difficult due the extensive genetic heterogeneity. The clade 131K has accumulated the same number of epitope mutations as 171K but has fewer non-epitope mutations relative to A/Texas/50/2012.

**Figure 6.**
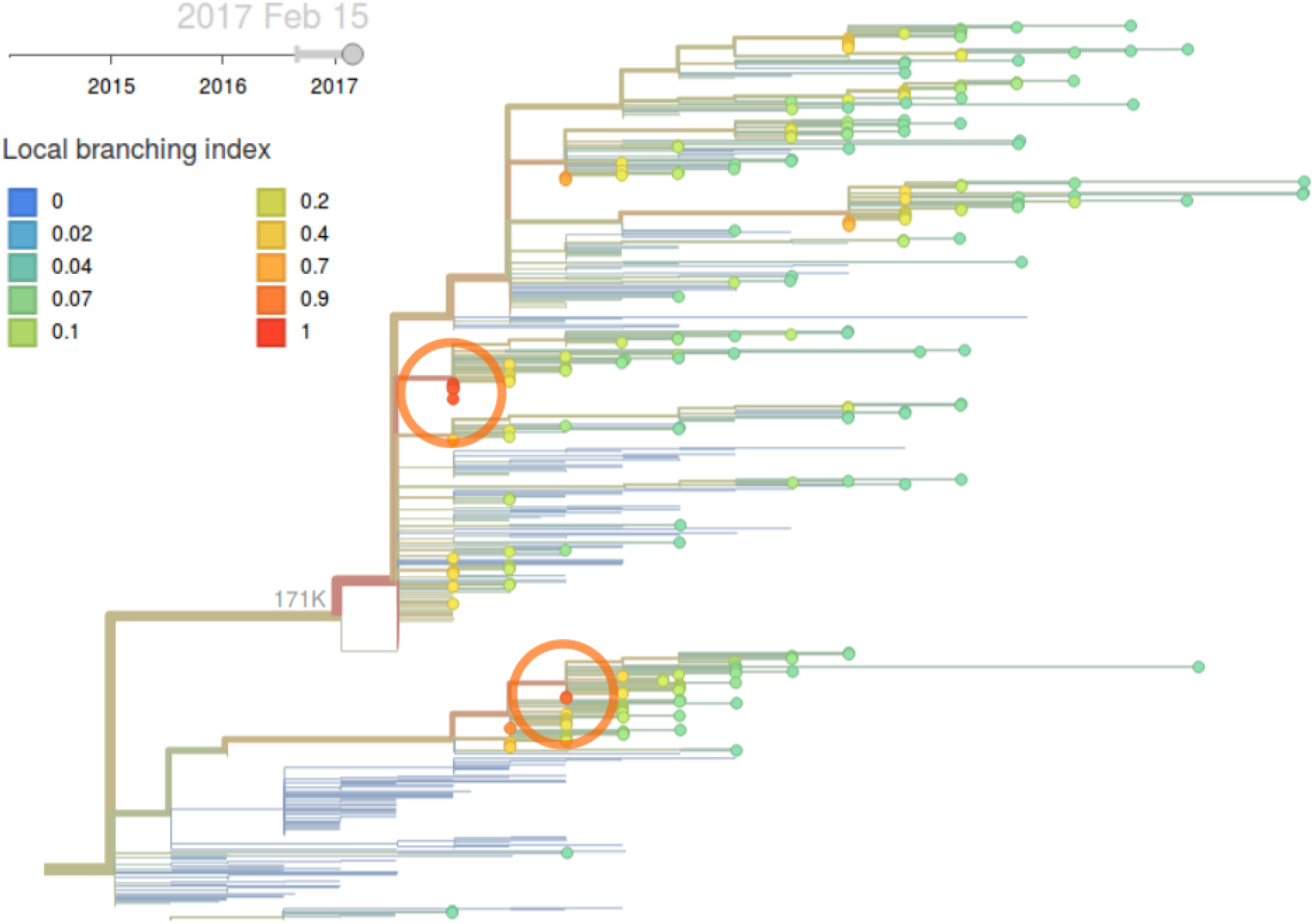
H3N2 phylogeny colored by local branching index.

*Given current patterns of clade growth and decline, we predict clades 171K/121K and 131K/142K to be the most successful of currently circulating clades. We expect both to increase in the coming months. Longer term projections are difficult. 171K/121K may continue to dominate, 131K/142K may displace 171K/121K or another mutation may appear that determines the eventual outcome.*

## A/H1N1pdm

**Very few H1N1pdm viruses have been observed in recent monthes. The dominant clade continues to be 6b.1 and there is little amino acid sequence variation within HA. The only notable subclade that has been growing recently is the clade bearing HA1:R205K/S183P. This clade is dominated by North American viruses and we see no evidence that this clade has a particular competitive advantage.**

As above, we base our primary analysis on a set of viruses collected between Jan 2015 and Feb 2017, comprising approximately 100 viruses per month where available and seeking to equilibrate sample counts geographically where possible (Fig. 7). However, in the case of H1N1pdm we were not able to collect a full 100 samples per month from the database in many months due to low absolute prevalence.

**Figure 7.**
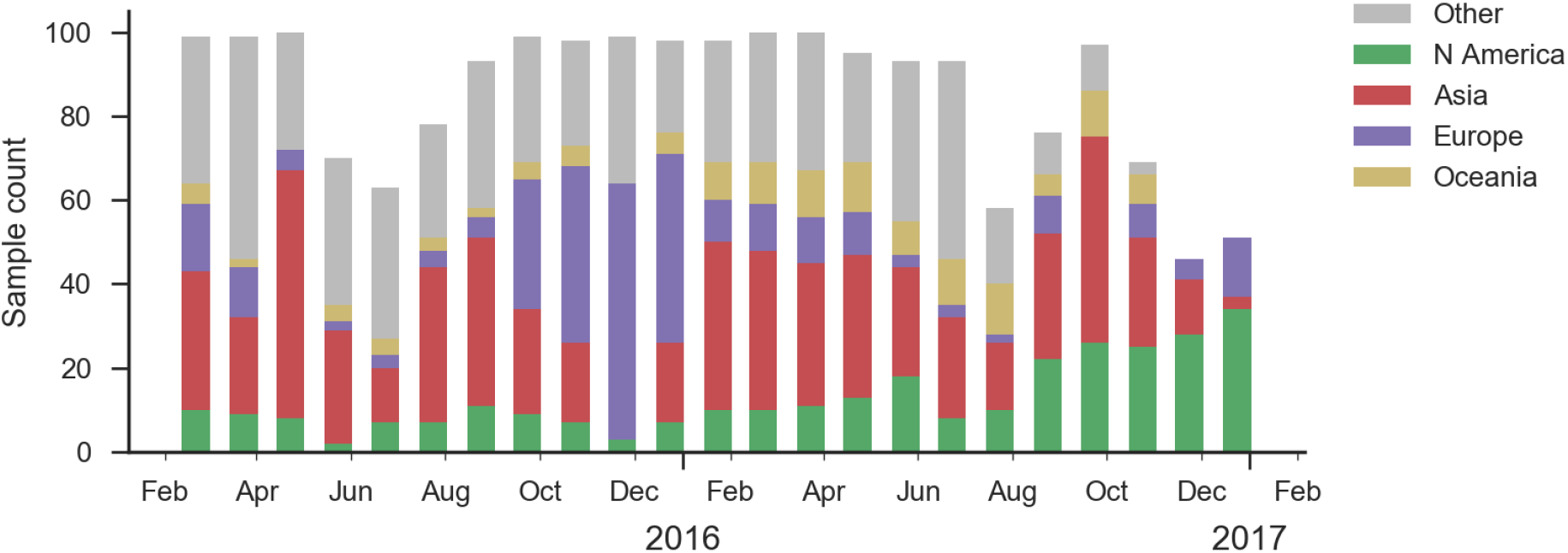
Sample counts through time and across regions. This is a stacked bar plot, so that in good months there are ~100 total samples and ~15 samples each from North America and from Europe.

There has been continued dominance of clade 6b.1 (characterized by mutation 162N) over clade 6b.2 (characterized by mutation 152T) (Fig. 8).

**Figure 8.**
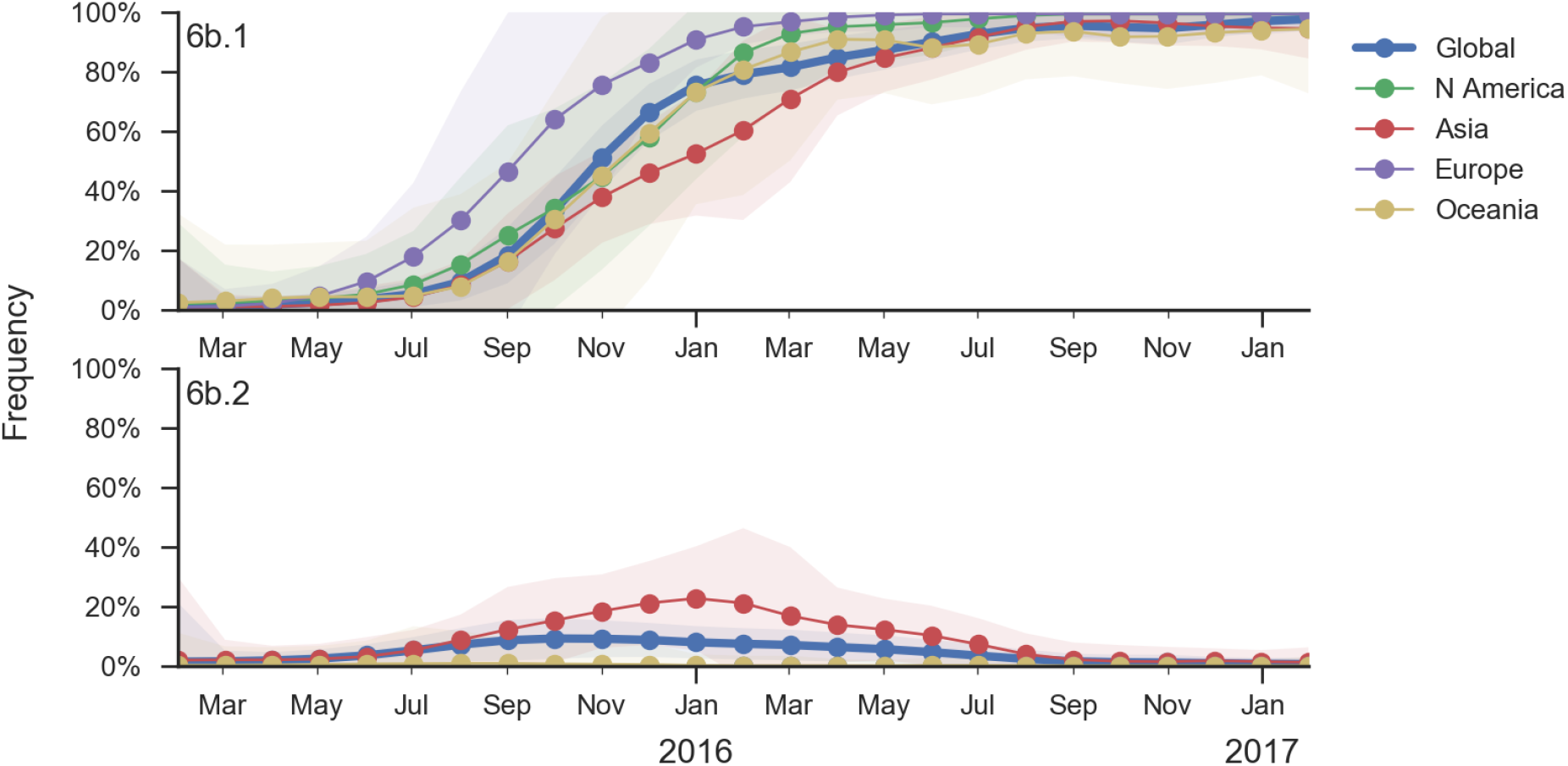
Frequency trajectories of H1N1pdm clades. We estimate frequencies of different clades based on sample counts and collection dates. We use a Brownian motion process prior to smooth frequencies from month-to-month. Transparent bands show an estimate the 95% confidence interval based on sample counts. The final point represents our frequency estimate for Feb 1 2017.

Within clade 6b.1 there is very little amino acid sequence variation (Fig. 9). The only mutations of any size are I166V, S183P and R205K. Interestingly, these three amino acid mutations occur in rapid succession along a single lineage, resulting in a series of nested clades.

**Figure 9.**
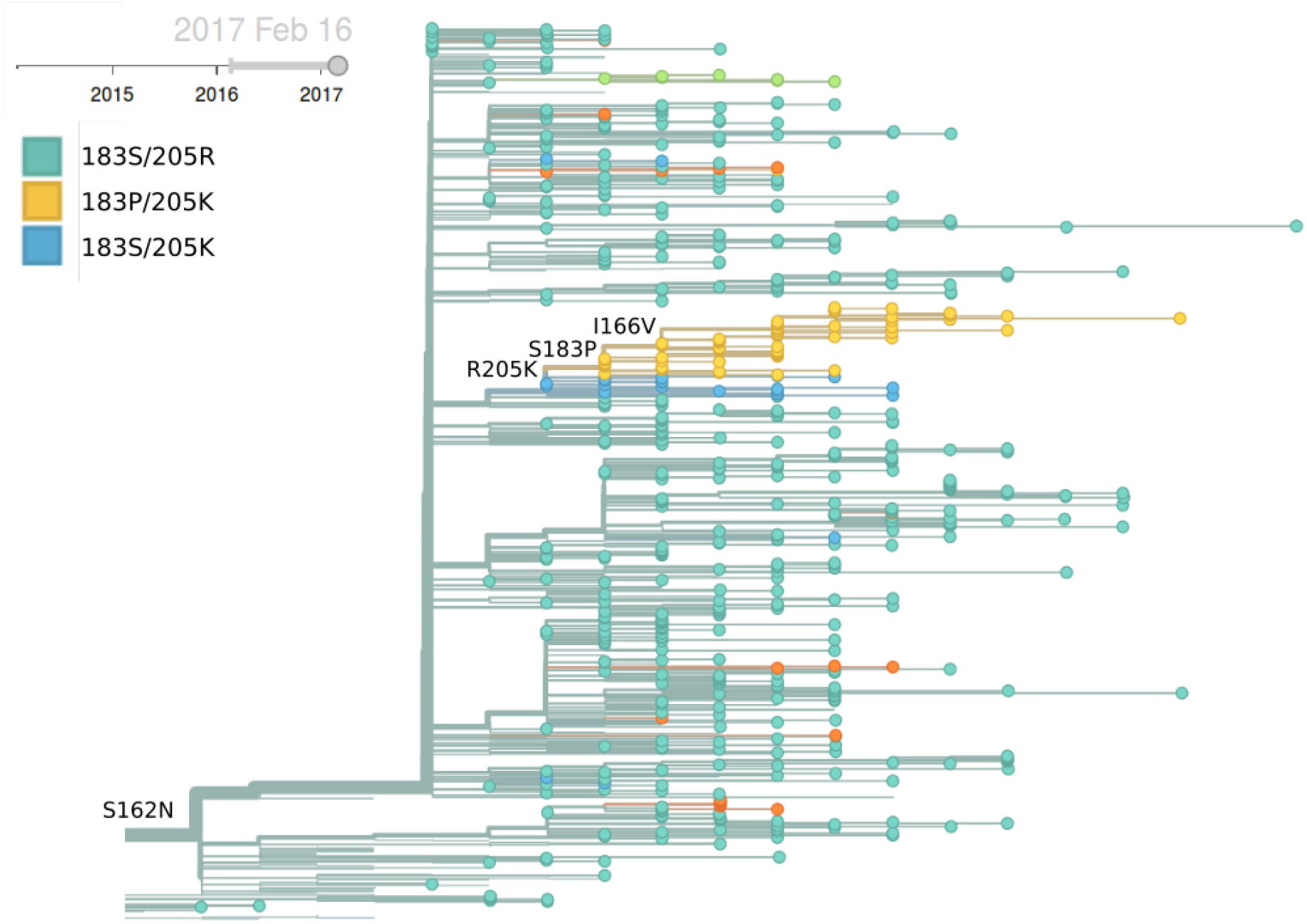
H1N1pdm / 6b.1 phylogeny colored by genotype.

Mutations 183P and 205K has been rising in Europe and North America starting in early-mid 2016, but this rise corresponds to a small number of cases and might turn out insignificant during the next H1N1pdm dominated season (Fig. 10). The pace of increase is only moderate, suggesting a lack of strong selective pressure. Additionally, no consistent antigenic variation is observed among recently circulating H1N1pdm viruses (according to primary infection in naive animals).

**Figure 10.**
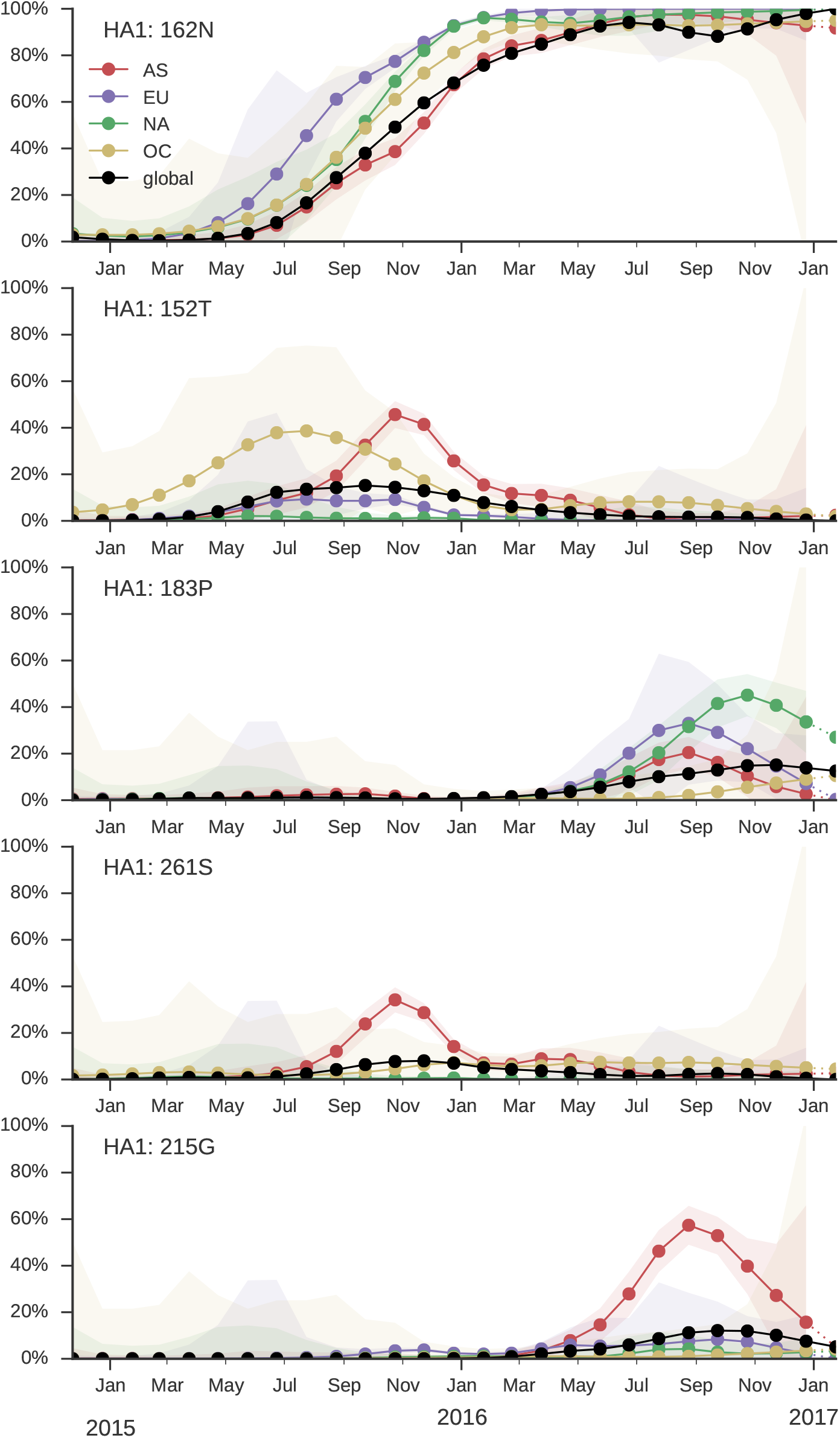
Frequency trajectories of 6b.1 subclades. We estimate frequencies of different clades based on sample counts and collection dates. We use a Brownian motion process prior to smooth frequencies from month-to-month. Transparent bands show an estimate the 95% confidence interval based on sample counts. The final point represents our frequency estimate for Feb 1 2017.

*Given the lack of competition, we predict the continued dominance of 6b.1 viruses.*

## B/Vic

**Clade 1A has continued to dominate and mutation 117V has all but taken over the global population. The rise of this mutation was fairly gradual and we have no evidence that it is associated with antigenic change or other benefit to the virus.**

As above, we base our primary analysis on a set of viruses collected between Jan 2015 and Feb 2017, comprising approximately 100 viruses per month where available and seeking to equilibrate sample counts geographically where possible (Fig. 11).

**Figure 11.**
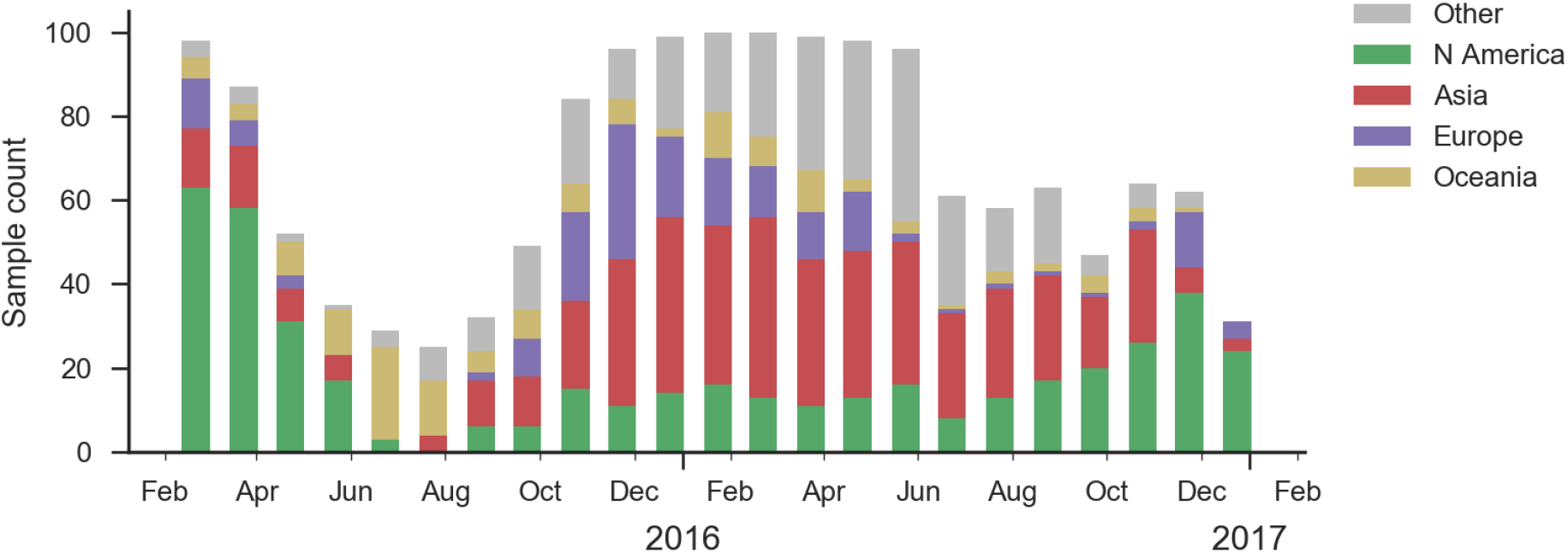
Sample counts through time and across regions. This is a stacked bar plot, so that in good months there are ^100 total samples and ~15 samples each from North America and from Europe.

We observe the continued dominance of the clade bearing mutations N129D, V146I and I117V (Fig. 12). There is very little HA amino acid variation within the 117V viruses with no subclades rising to appreciable frequency. There is deletion at AA site 162 that first appeared in mid-2016, but remains at low global frequency.

**Figure 12.**
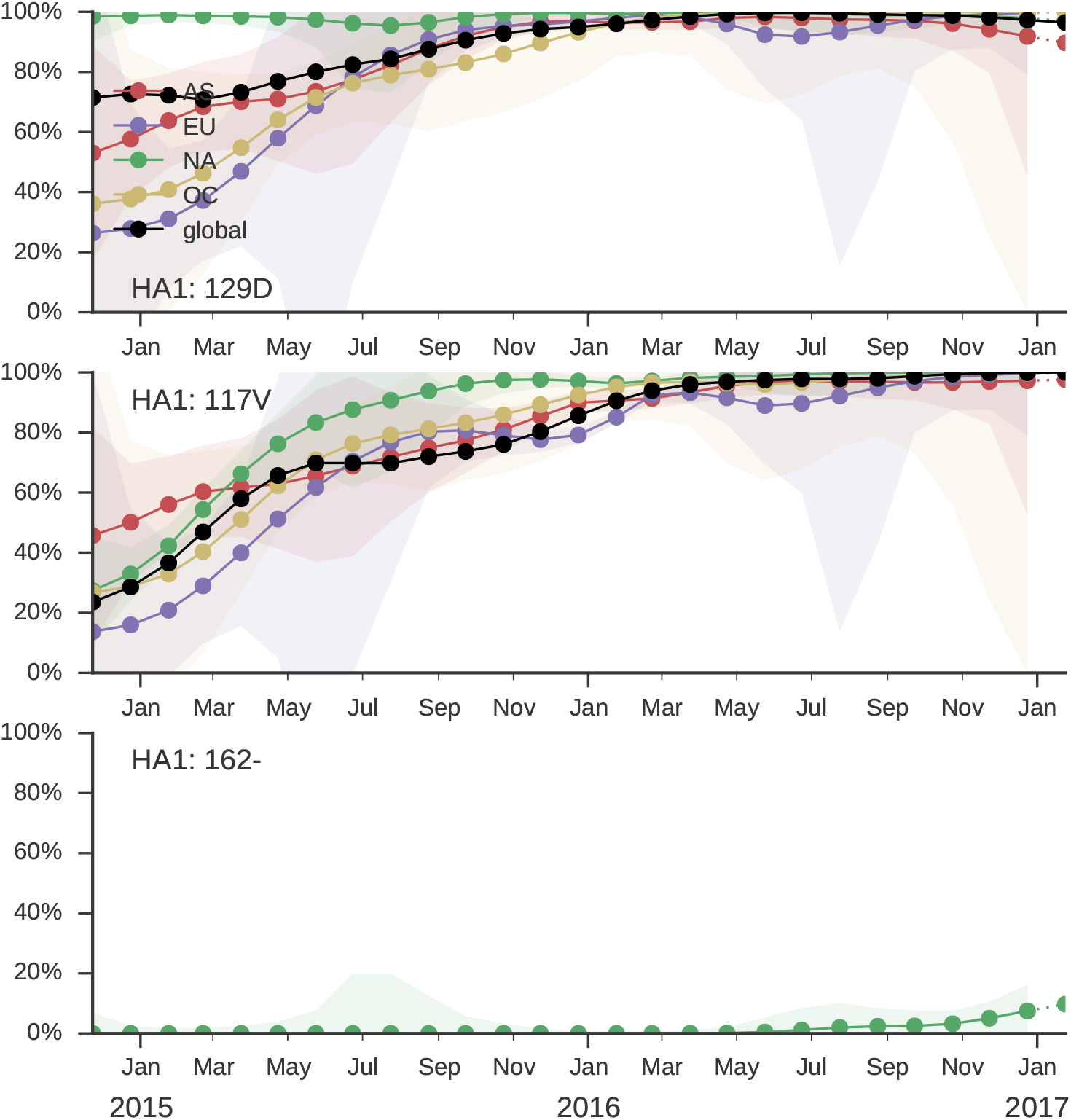
Frequency trajectories of B/Vic variants. We estimate frequencies of different clades based on sample counts and collection dates. We use a Brownian motion process prior to smooth frequencies from month-to-month. Transparent bands show an estimate the 95% confidence interval based on sample counts. The final point represents our frequency estimate for Feb 1 2017.

Neither the local branching index nor clade frequencies single out a particular variant within the 1A/I117V clade (Fig. 13).

**Figure 13.**
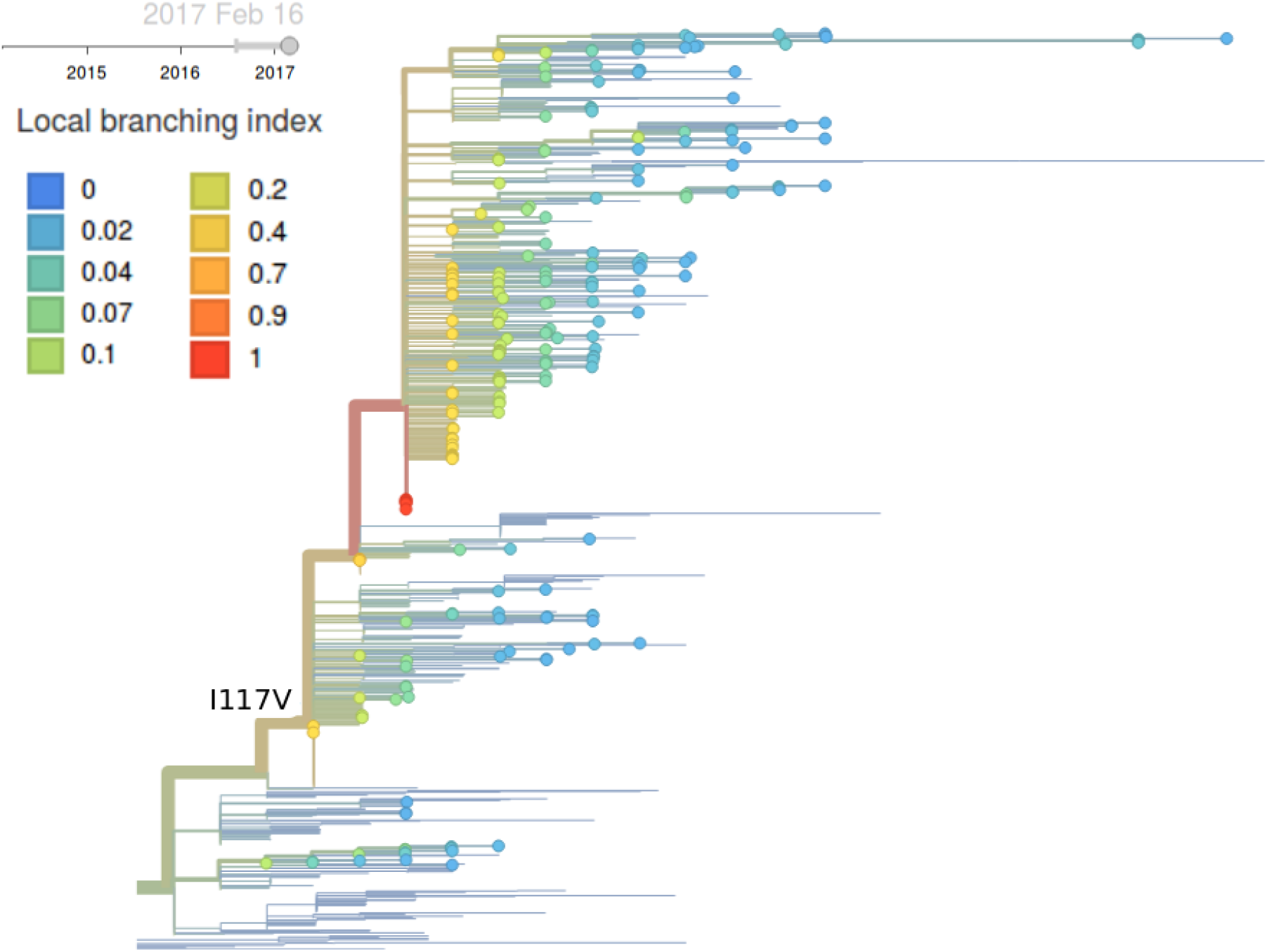
B/Vic phylogeny colored by local branching index.

*Given the lack of competition, we predict the continued dominance of 117V viruses.*

## B/Yam

**Clade 3 has continued to dominate. Within clade 3, a clade with mutation HA1:251V is globally at frequency of about 80% throughout 2016. Within this clade, mutation 211R is at 25% frequency. In addition, a clade without prominent amino acid mutations has been rising throughout 2016.**

As above, we base our primary analysis on a set of viruses collected between Jan 2015 and Feb 2017, comprising approximately 100 viruses per month where available and seeking to equilibrate sample counts geographically where possible (Fig. 14).

**Figure 14.**
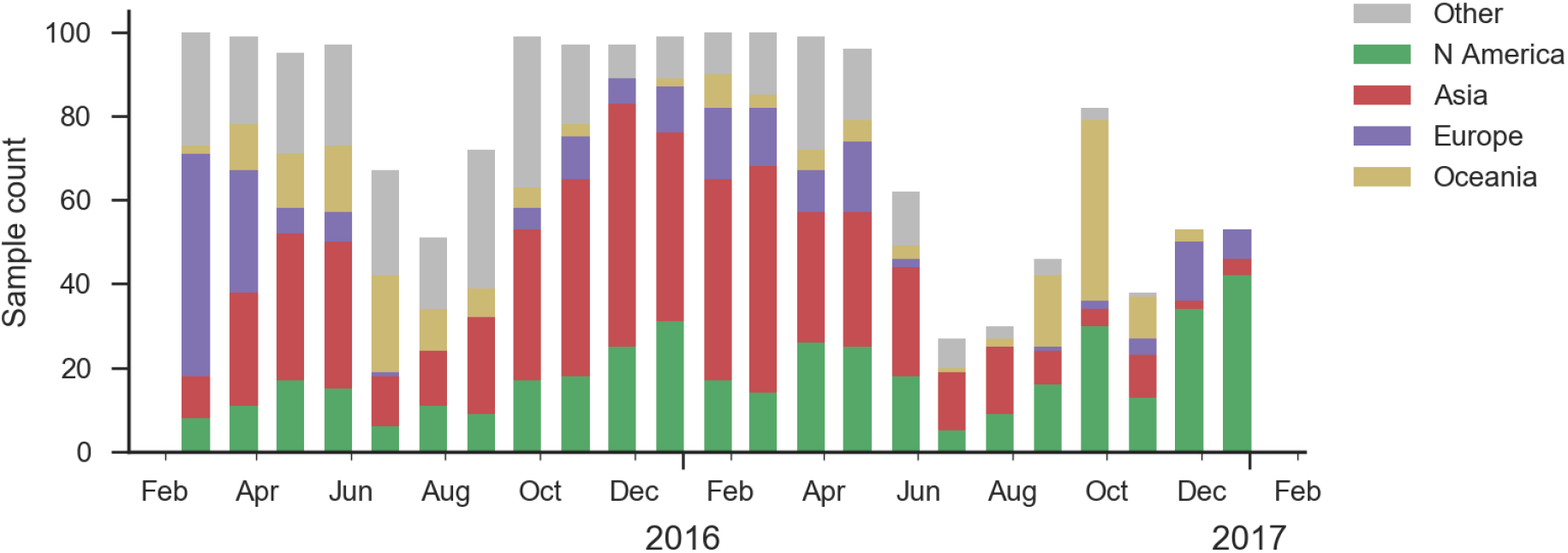
Sample counts through time and across regions. This is a stacked bar plot, so that in good months there are ~100 total samples and ~15 samples each from North America and from Europe.

Clade 3 viruses have continued to dominate with clade 2 viruses scarcely observed throughout 2016. Within clade 3, mutations L172Q and M251V remain dominant and mutation K211R continues to persist at low frequency (Fig. 15, 16). There is little evidence of emerging clades of strong selective effect.

**Figure 15.**
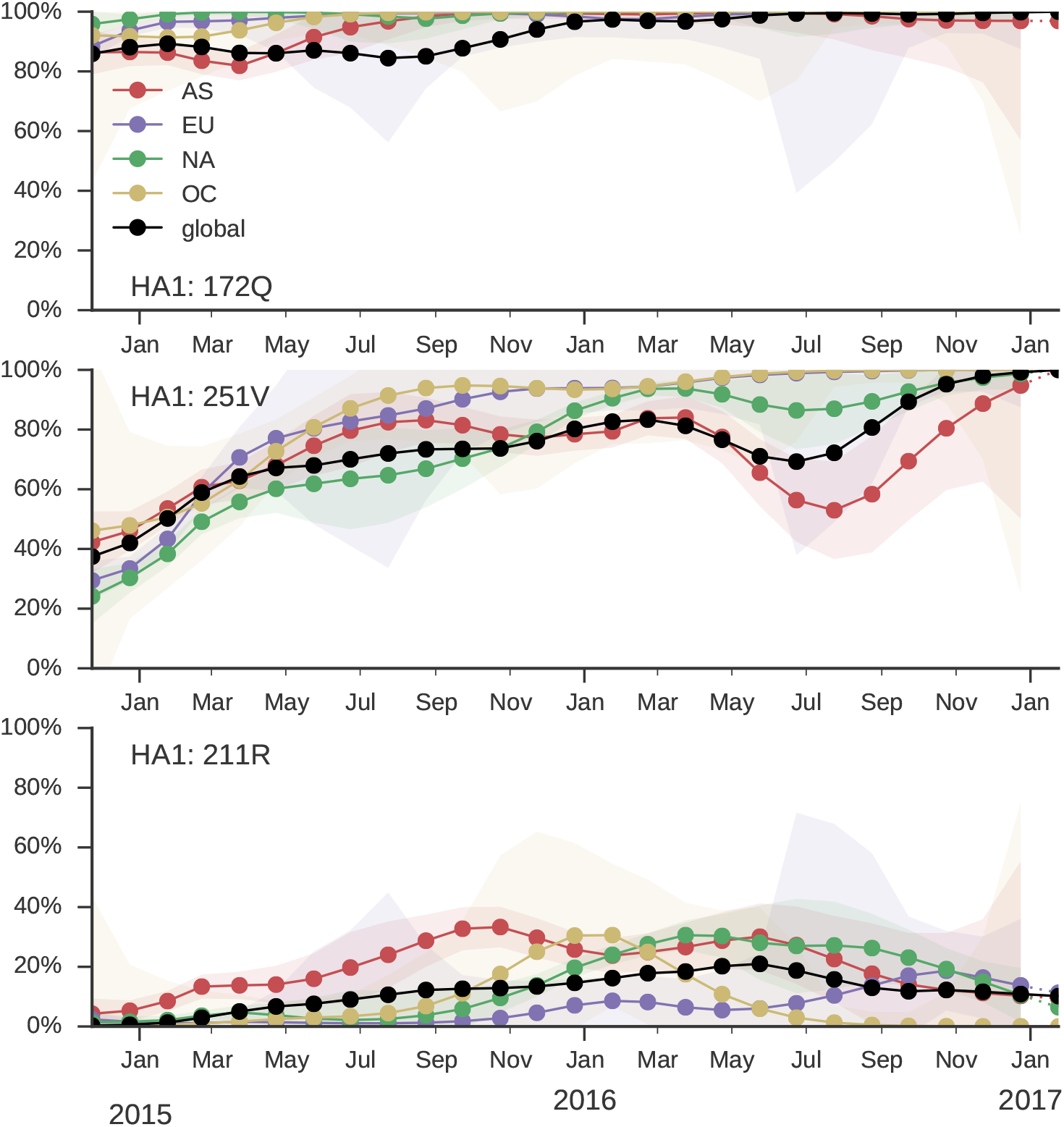
Frequency trajectories of B/Yam variants. We estimate frequencies of different clades based on sample counts and collection dates. We use a Brownian motion process prior to smooth frequencies from month-to-month. Transparent bands show an estimate the 95% confidence interval based on sample counts. The final point represents our frequency estimate for Feb 1 2017.

**Figure 16.**
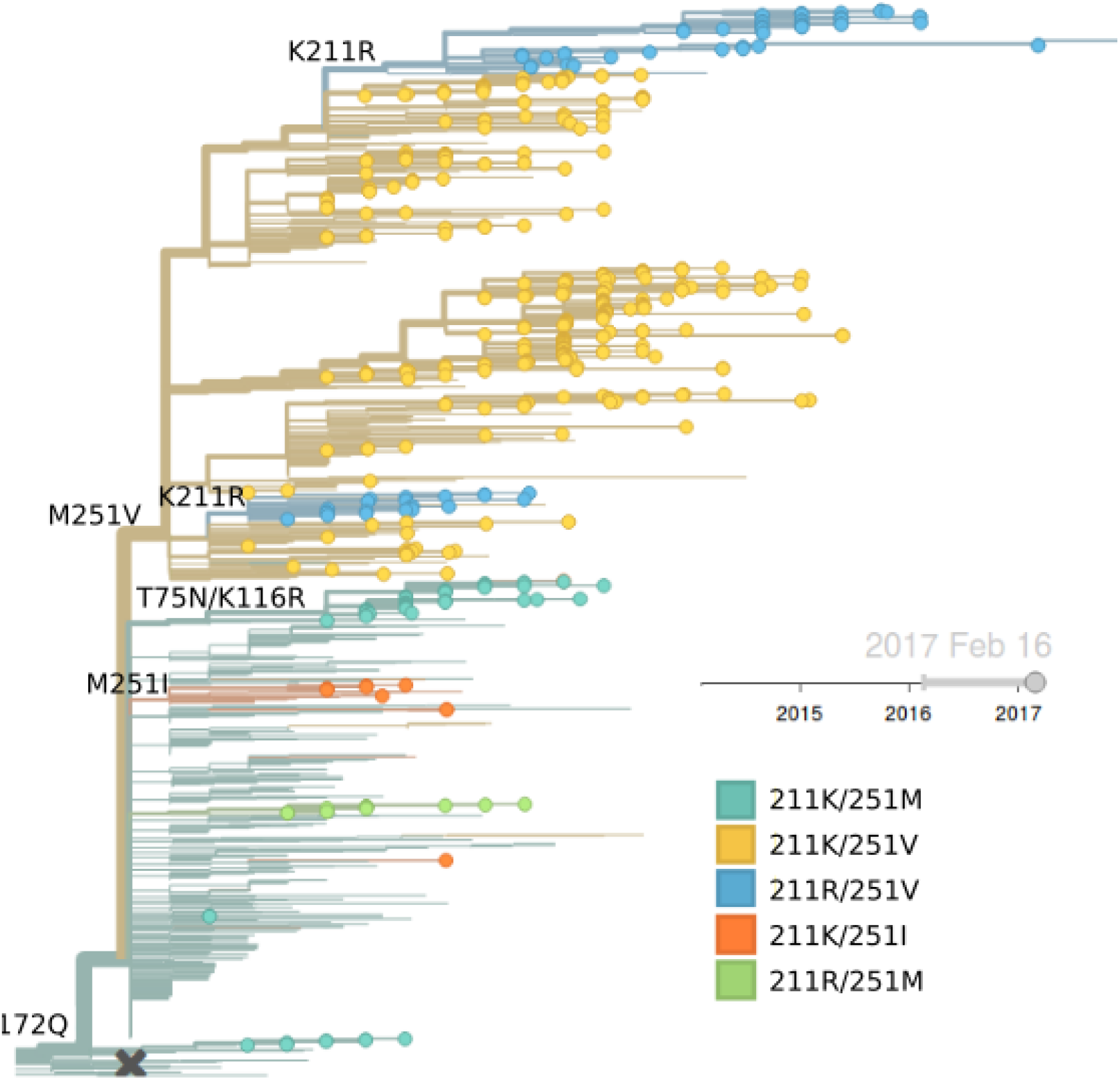
B/Yam phylogeny colored by genotype.

The LBI highlights the central clade in the tree below (Fig. 17). This clade has no amino acid mutations in HA along its backbone relative to its parent clade 251V.

**Figure 17.**
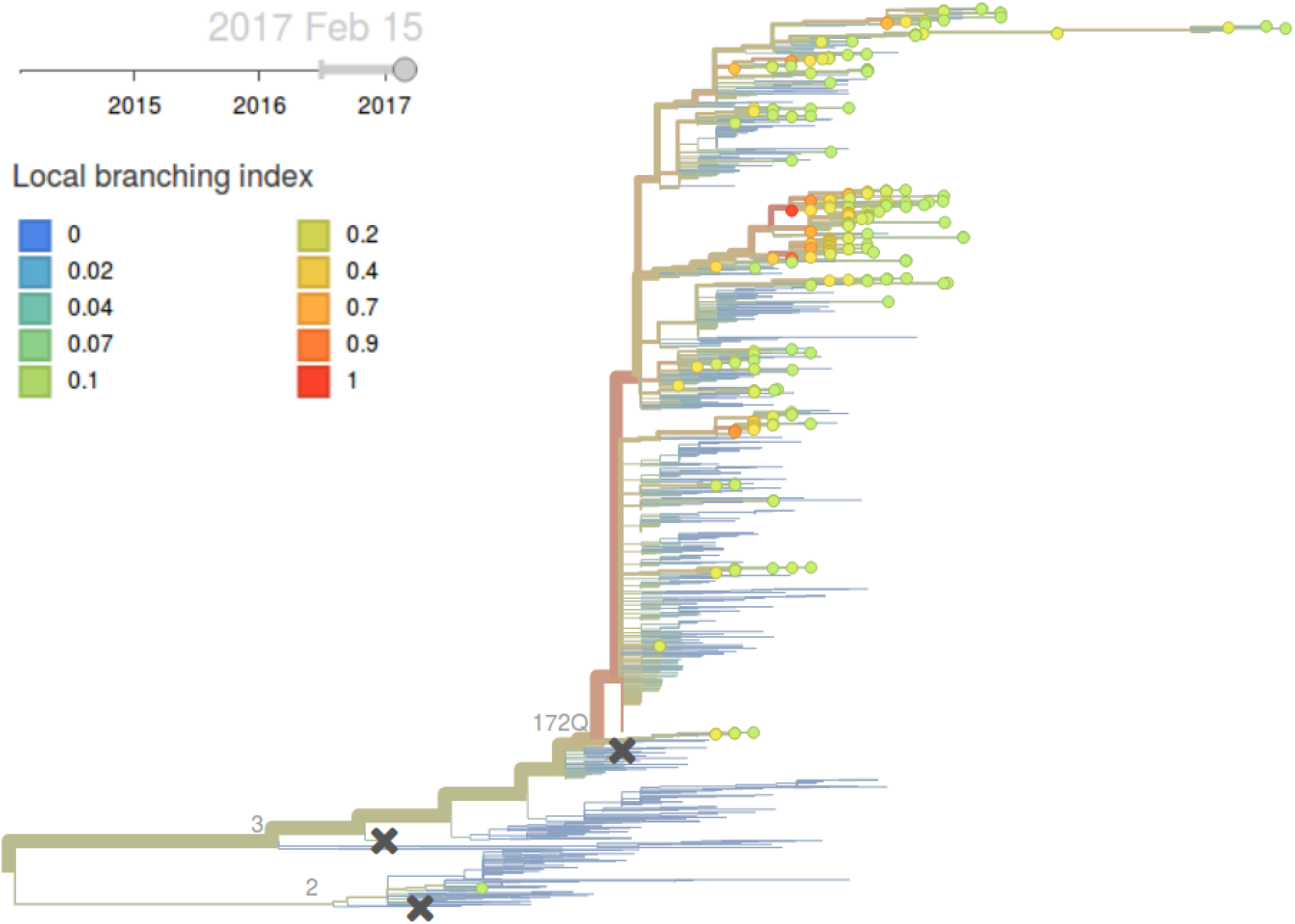
B/Vic phylogeny colored by local branching index.

*Genetic variation within 172Q/251V viruses is beginning to develop, but given lack of credible competition, we expect 172Q/251V to continual to dominate in the coming months.*

